# Evaluation of Antimicrobial, Antioxidant, Total Phenolic, Total Flavonoids, Metal Content and Proximate Potential of *Solanum xanthocarpum* L. (Solanaceae)

**DOI:** 10.1101/2020.02.20.957381

**Authors:** Nosheen Azhar, Muhammad Qayyum Khan, Asia Bibi, Muhammad Shoaib, Sadia Mumtaz, Tahira Batool, Saima Ashraf, Shabir Ijaz, Sadiqa Firdous, Faryal Sharif

## Abstract

The current research work is focused on screening of antimicrobial, antioxidant, total phenolic, total flavonoid, metal estimation and proximate potential of four parts (fruits, leaves, stem bark and root bark) of *S. xanthocarpum*. Antimicrobial potential of dried crude extracts of *S. xanthocarpum* were evaluated against two gram positive bacteria (*S. aureus, P. vulgaris*), three gram negative bacteria (*P. aeruginosa, K. pneumonia, E. coli*), three fungi (*A. flavus, F. solani, stolonifer*) and two yeasts (*S. cerevisiae, C. albicans*) by using disc diffusion assay. Five organic solvents ranging from non-polar to highly polar were used for extraction of active metabolites. Amongst all the parts of *S. xanthocarpum* tested, antimicrobial activity of methanol extract of fruits (14.67±0.33) against *P. aeruginosa*, ethanolic extract of leaves against *P. vulgaris* (14±0.58), stem bark methanolic extract against *S. aureus* (13±0.58) and stem bark methanolic extract against *P. vulgaris* (17.67±0.33) were found to be more significant. All other extracts also showed promising antimicrobial activity against bacterial pathogens. Among fungal pathogens, *R. stolonifer* and *S. cerevisiae* were found to be more sensitive to extracts of *S. xanthocarpum*. Gram negative bacteria exhibited more resistance than gram positive bacteria. However, fungi were found to be more resistant than bacteria. All the extracts showed antioxidant activity. However, methanol extract of stem bark of *S. xanthocarpum* with IC_50_ value of 0.323102 mg mL^−1^ showed maximum antioxidant potential. Total phenolic contents ranged from 12.3541±1.73 to 23.2942±1.33 Pmol GA/ug. However, highest flavonoid content was found in the stem bark extract (17.8480±1.75 ugRutin/ug) and lowest in leaves extract of *S. xanthocarpum* (2.4806±0.59 ugRutin/ug). Total metal contamination in four parts of *Solanum xanthocarpum* (fruits, leaves, stem bark and root bark) was estimated by atomic absorption spectroscopy and results showed Cadmium contamination in its stem bark and root bark, Chromium contamination in leaves, stem bark and root bark, Copper and Magnesium contamination in all parts of *S. xanthocarpum* and Maganese contamination in leaves critically above the standard permissible limits. The proximate analysis of the *Solanum xanthocarpum* revealed that stem bark is a poor source of lipid (3.42%) and high carbohydrate (50.07%) and ash (16.50%) contents. Whereas, root bark has highest wet moisture content (65%), dry moisture content (20%) and lowest fiber (13%). Highest energy (285.455%), protein (8.32%), fat (9.79%) and lowest amount of ash (13%) was found in fruits. This composition shows that the *Solanum xanthocarpum* could be considered as a good source of carbohydrate, moisture and energy. These results revealed that over all, methanolic extract of the *S. xanthocarpum* is richest in phenolic, flavonoid and nutritional contents as well as most potent against bacterial and fungal pathogens. Therefore, further investigation is recommended to isolate, screen and characterize their active metabolites.

## Introduction

Medicinal plants contain superlative nutritional composition and phytochemicals which are used in various biological functions and treatment of various diseases in the human body [1]. It has been observed that in majority of the developing countries, medicinal plants are widely used for disease prevention, health maintenance and treatment of various mental and physical illnesses [2]. Pakistan is naturally blessed with diverse climatic condition from Northern Himalayian, central hot region to southern moderate conditions which create and facilitate rich and diverse metabolic engineering of therapeutic natural products in its medicinal flora. Additionally, Azad Kashmir, Pakistan has rich culture and knowledge of using these medicinal species to treat and prevent various ailments. Biological screening of these medicinal plants is the starting point to the discovery of medicinal natural products from medicinal plants. In Azad Kashmir, medicinal plants are readily available and cheapest sources of phytochemicals and biological contents [3] and therefore could also benefit the populace with their medicinal properties.

*Solanum xanthocarpum* Schrad. and Wendl. belongs to family Solanaceae is a very prickly diffuse bright-green perennial herb, somewhat woody at the base [4]. *S. xanthocarpum* has been reported as safe for human consumption due to its non-toxicity [5]. *Solanum xanthocarpum* has profound use in Ayurveda and folklore medicine. The whole plant is useful in vitiated conditions of lumbago, helminthiasis, anorexia, epilepsy, inflammations, dental claries, constipation, dyspepsia, leprosy, skin diseases, hypertension, flatulence, cough, asthma, bronchitis, fever, hiccough, haemorrhoids, vata and kapha. Importantly, *Solanum xanthocarpum* is the superlative precursor for the discovery and development of molecular pharmaceuticals. Therefore, present study was designed and conducted to evaluate antimicrobial, antioxidant, total phenolic, total flavonoids, metal content and proximate potential of *Solanum xanthocarpum* L.

## Material and methods

### Plant collection and processing

*S. xanthocarpum* (fruits, leaves, stem bark and root bark) locally known as Mohkrri were collected from Rayala, Dhirkot Azad Kashmir and authenticated. Each part of the plant was washed properly and shade dried for 10 days at room temperature followed by grinding to fine powder. The samples were stored at 4°C till further studies. For preparation of sample solutions, 100 g dried powder of each part of plant was separately macerated in 300 mL of petroleum ether, chloroform, acetone, ethanol, methanol and kept at room temperature (25±2°C) for 10 days. After maceration period, the samples were filtered and extracts were dried on rotary evaporator at 60°C and reduced pressure.

### Preparation of extract for antimicrobial activity

10 mg of each dried crude extract of fruits, leaves, stem bark and root bark were dissolved in 10 mL of respective solvents i.e. petroleum ether, chloroform, acetone, ethanol and methanol in sterile eppendorf tubes.

### Pathogenic isolates

For antimicrobial activity, clinical pathogens including Gram Positive Bacteria (*Staphylococcus aureus*), Gram Negative Bacteria (*Escherichia coli, Klebsiella pneumoniae, Proteus vulgaris and Pseudomonas aeruginosa*), Fungi (*Aspergillus flavus, Fusarium solani* and *Rhizopus stolonifer*), Yeast (*Saccharomyces cerevisiae* and *Candida albicans*) were *obtained* from Microbiology Lab of National Institute of Health (NIH), Islamabad and Combined Military Hospital (CMH), Muzaffarabad.

### Antimicrobial assay

In vitro screening of crude extracts was carried out by disc diffusion method [6]. Antibacterial and antifungal potential of crude extracts was compared with ciprofloxacin, tetracycline, nystatin against clinical pathogens. The plates containing bacterial culture were incubated for 24 hours at 37°C. However, fungal and yeast dilutions were incubated at 25°C for 72 hours. After incubation, of the zone of inhibition was measured (mm). All the experiments were performed in triplicates.

### Antioxidant activity

The DPPH (2,2-diphenyl-1-picrylhydrazyl) radical scavenging assay was used [7] to measure antioxidant activity. Extracts with various concentrations (1 mg/ul, 2 mg/ul and 5 mg/ul) were used for assay. The inhibition percentage radicals scavenging activity was calculated by using following formula;

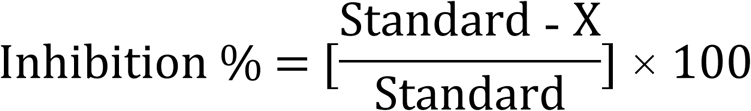

X= Sample-Blank

Where standard is the absorbance of control reaction (containing all reagents except the test compounds). 50 percent inhibition (IC_50_) of each extract concentrations against graph of inhibition was calculated by applying SSP10 software.

### Total phenolic and flavonoid content

Total phenolic content in methanolic extracts of all parts of plant sample was determined by Folin-Ciocalteu method [8] and expressed as gallic acid equivalents (Pmol GA/ µg). Total flavonoid content in methanolic extracts of all parts of plant sample was measured as described previously by [9] and was calculated as rutin equivalents (µg Rutin/g).

### Sample preparation for the estimation of metal content

Powdered plant material (1 gm) was taken in a glass beaker (Pyrex) and (2:1 V/V) of concentrated nitric acid (6 mL) and hydrogen peroxide (3 mL) were added into the beaker and mixture was stirred slowly at room temperature followed by heating at 70°C for three hours. Small volumes of (HNO_3_) and H_2_O_2_ were added to this mixture along with continuous heating at 80°C till a clear solution was obtained. Futher heated until a semi dry mass was obtained. The semi dry mass was solubilized in 10 mL HNO_3_ (0.1 M) and was transferred to an Erlenmeyer flask and final volume was adjusted to 50 mL with double distilled water. Analysis of trace elements was carried out by Atomic Absorption Spectrophotometer (AAS, AAnalyst700) using respective wavelength in the range of 190-900 nm. Total volume of solution was made 50 mL by adding double distilled water. To calculate the mean of three readings winlab 32 program was used.

### Proximate analysis

The parameters determined for proximate analyses include moisture content (wet, dry), ash, fiber, fat, crude protein, carbohydrate and energy. The parameters for proximate analysis of four parts of *S. xanthocarpum* were measured in percentage following the protocol by [10, 11] with slight modification.

### Statistical analysis

All values were expressed as mean ± standard error. By using one way analysis of variance (ANOVA) technique, the data was analyzed [12]. The mean values were compared by using LSD at 5% (0.05) probability level by using SPSS 13.0 software.

## Results and discussion

### Antimicrobial activities of different parts of *S. xanthocarpum*

Different parts of *S. xanthocarpum* showed promising antimicrobial potential against selected clinical isolates. Table 1 and Fig 1 represents the results of fruit of *S. xanthocarpum*. Results of the sensitivity tests were expressed as (0) for no sensitivity, (7-10 mm) for low sensitivity, (11-18 mm) for moderate sensitivity and (> 19 mm) for high sensitivity. It is evident from the results that the methanolic fruit extract of *S. xanthocarpum* showed prominent activity against *P. aeruginosa* (14.67±0.33 mm) and moderate activity against *S. aureus* (13.33±0.33 mm) and *E. coli* (12.33±0.33 mm) respectively. The zone of inhibition of methanolic extract (13.33±0.67d) was found close to tetracycline (14.33±0.33) against *S.aureus*. Whereas, it showed marginal activity against *R. stolonifer* (9.67±0.33 mm) and *P. vulgaris* (8.67±0.33 mm). Present findings are in line to the previous report which suggested that methanol extract showed higher potential [13]. All other extracts were also found to possess considerable inhibitory activity. However, most of the fungi and yeasts were found to be resistant to fruit extract.

**Table 1:** Antimicrobial activity of *S. xanthocarpum* fruit extract against clinical isolates.

| Mean diameter of zones of inhibition (mm) ± Standard Error of Means (S.E.M) |  |  |  |  |  |  |  |  |  |
| --- | --- | --- | --- | --- | --- | --- | --- | --- | --- |
| Sr. No | Microorganisms | Petroleum ether | Chloroform | Acetone | Ethanol | Methanol | Tetracycline | Ciprofloxacin | Nystatin |
| 1 | *S. aureus* | 8.67±0.33e | 9.33±0.33e | 12.67±0.33e | 10.67±0.33 | 13.33±0.67d | 14.33±0.33d | 33.33±0.33a | --- |
| 2 | *P. aeruginosa* | 8.33±0.33e | 13.67±0.33e | 9.67±0.33e | 11.67±0.33e | 14.67±0.33d | 20.33±0.33c | 33.33±0.33a | --- |
| 3 | *K. pneumonia* | 7.67±0.33g | 7.67±0.33g | 11.67±0.33d | 10.67±0.33e | 10.67±0.33e | 15.33±0.33d | 29.67±0.33b | --- |
| 4 | *P. vulgaris* | 12.33±0.67d | 8.67±0.33e | 13.67±0.33e | 12±0.58e | 9.67±0.33e | 19.67±0.33e | 32.33±0.67a | --- |
| 5 | *E. coli* | 9.67±0.33f | 9.33±0.33f | 11.67±0.33e | 11.33±0.33e | 12.33±0.33e | 17.67±0.33d | 20.67±0.33c | --- |
| 6 | *A. flavus* | 0.00±0.00h | 0.00±0.00h | 0.00±0.00h | 0.00±0.00h | 0.00±0.00h | --- | --- | 20.67±0.33d |
| 7 | *F. solani* | 0.00±0.00h | 0.00±0.00h | 0.00±0.00h | 0.00±0.00h | 0.00±0.00h | --- | --- | 28.33±0.33b |
| 8 | *R. stolonifer* | 8.33±0.33g | 12.67±0.33f | 11.67±0.33f | 0.00±0.00h | 9.67±0.33 | --- | --- | 39.67±0.33a |
| 9 | *S. cerevisiae* | 0.00±0.00h | 0.00±0.00h | 11.67±0.33f | 0.00±0.00h | 0.00±0.00h | --- | --- | 23.33±0.33c |
| 10 | *C. albicans* | 0.00±0.00h | 0.00±0.00h | 0.00±0.00h | 0.00±0.00h | 0.00±0.00h | --- | --- | 17.67±0.33e |

Note: The means followed by same letters are non-significant

**Figure 1.**
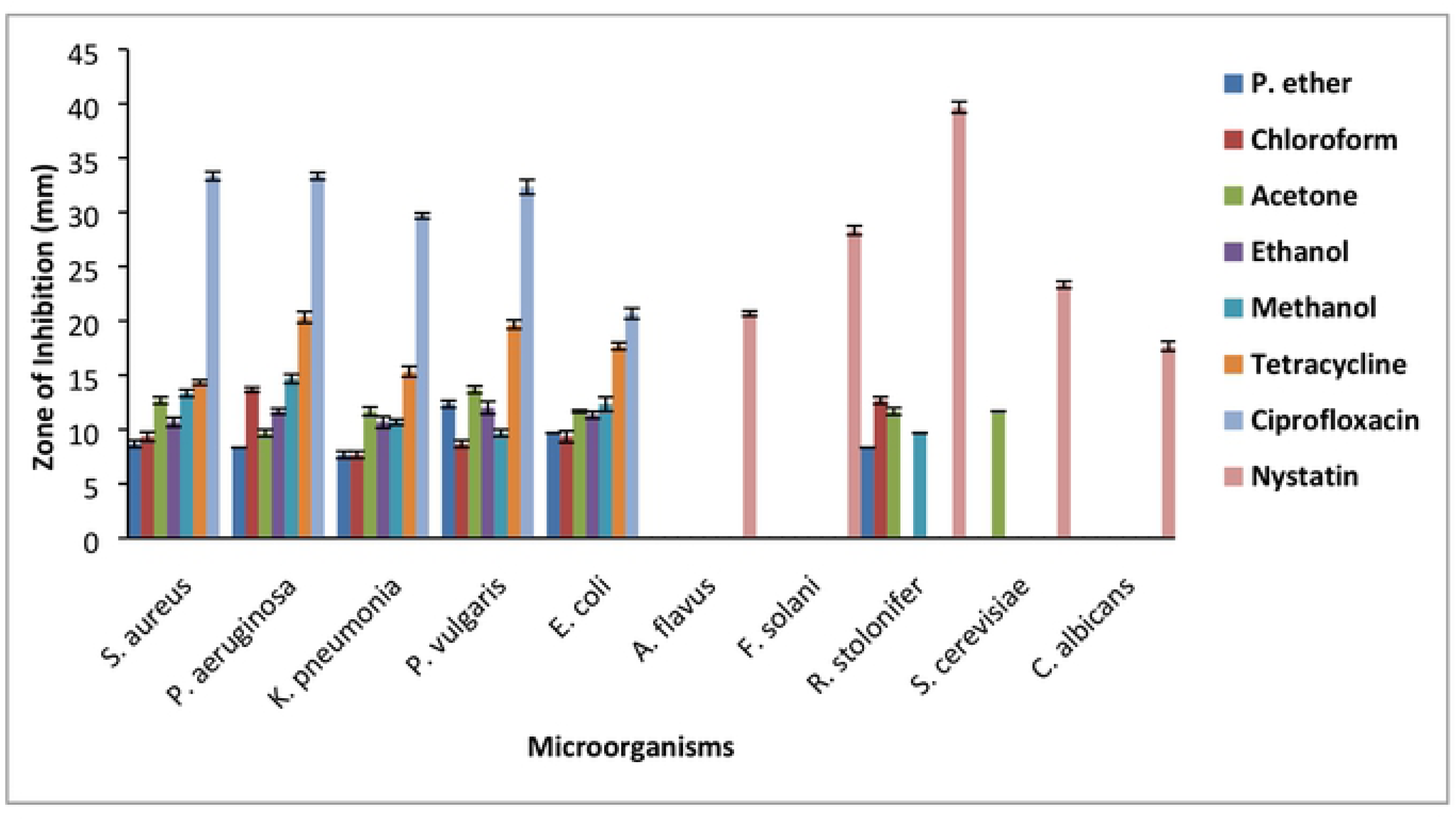
Antimicrobial activities of fruits extract of *S.Xanthocarpum*.

The ethanol extract of leaves showed highest antimicrobial activity (14±0.58 mm) against *P. vulgaris*. The methanolic extract of leaves of selected plants also showed maximum activity against *P. aeruginosa* (13.67±0.33 mm) (Table 2 and Fig 2). These results are well correlated with previous studies [14] in which ethanol extract showed higher activity. All the bacteria were found sensitive against all the extracts. Among fungi, only *R. stolonifer* showed sensitivity towards leaf extracts of acetone and chloroform with zone of inhibition of 11.33±0.33 mm and 8.33±0.33 mm respectively. Similar to the present findings, [15] reported superior antibacterial activity of *Bellis prennis extract than antifungal*.

**Table 2:** Anti-microbial activity of *S. xanthocarpum* leaves extract against clinical isolates.

| Mean diameter of zones of inhibition (mm) ± Standard Error of Means (S.E.M) | | | | | | | | | |
| --- | --- | --- | --- | --- | --- | --- | --- | --- | --- |
| S. No | Microorganisms | Petroleum ether | Chloroform | Acetone | Ethanol | Methanol | Tetracycline | Ciprofloxacin | Nystatin |
| 1 | *S. aureus* | $7.67 \pm 0.33$ | $10.33 \pm 0.33e$ | $10.67 \pm 0.33e$ | $10.67 \pm 0.33e$ | $11.33 \pm 0.33e$ | $14.33 \pm 0.33d$ | $33.33 \pm 0.33a$ | --- |
| 2 | *P. aeruginosa* | $7.33 \pm 0.00f$ | $13.67 \pm 0.33d$ | $9.67 \pm 0.33e$ | $13.33 \pm 0.67d$ | $13.67 \pm 0.33d$ | $20.33 \pm 0.33c$ | $33.33 \pm 0.33a$ | --- |
| 3 | *K. pneumonia* | $12.33 \pm 0.33e$ | $7.33 \pm 0.33f$ | $11.67 \pm 0.33e$ | $10.67 \pm 0.33e$ | $9.67 \pm 0.33e$ | $15.33 \pm 0.33d$ | $29.67 \pm 0.33b$ | --- |
| 4 | *P. vulgaris* | $11.33 \pm 0.33e$ | $8.33 \pm 0.33f$ | $13.67 \pm 0.33d$ | $14 \pm 0.58d$ | $8.67 \pm 0.33f$ | $19.67 \pm 0.33e$ | $32.33 \pm 0.67a$ | --- |
| 5 | *E. coli* | $9.67 \pm 0.00e$ | $11.33 \pm 0.33e$ | $12.67 \pm 0.33e$ | $12 \pm 0.33e$ | $12.33 \pm 0.33e$ | $17.67 \pm 0.33d$ | $20.67 \pm 0.33c$ | --- |
| 6 | *A. flavus* | $0.00 \pm 0.00h$ | $0.00 \pm 0.00h$ | $0.00 \pm 0.00h$ | $0.00 \pm 0.00h$ | $0.00 \pm 0.00h$ | --- | --- | $20.67 \pm 0.33d$ |
| 7 | *F. solani* | $0.00 \pm 0.00h$ | $0.00 \pm 0.00h$ | $0.00 \pm 0.00h$ | $0.00 \pm 0.00h$ | $0.00 \pm 0.00h$ | --- | --- | $28.33 \pm 0.33b$ |
| 8 | *R. stolonifer* | $0.00 \pm 0.00h$ | $8.33 \pm 0.33g$ | $11.33 \pm 0.33f$ | $0.00 \pm 0.00h$ | $0.00 \pm 0.00h$ | --- | --- | $39.67 \pm 0.33a$ |
| 9 | *S. cerevisiae* | $0.00 \pm 0.00h$ | $0.00 \pm 0.00h$ | $0.00 \pm 0.00h$ | $0.00 \pm 0.00h$ | $0.00 \pm 0.00h$ | --- | --- | $23.33 \pm 0.33c$ |
| 10 | *C. albicans* | $0.00 \pm 0.00h$ | $0.00 \pm 0.00h$ | $0.00 \pm 0.00h$ | $0.00 \pm 0.00h$ | $0.00 \pm 0.00h$ | --- | --- | $17.67 \pm 0.33e$ |

Note: The means followed by same letters are non-significant

**Figure 2.**
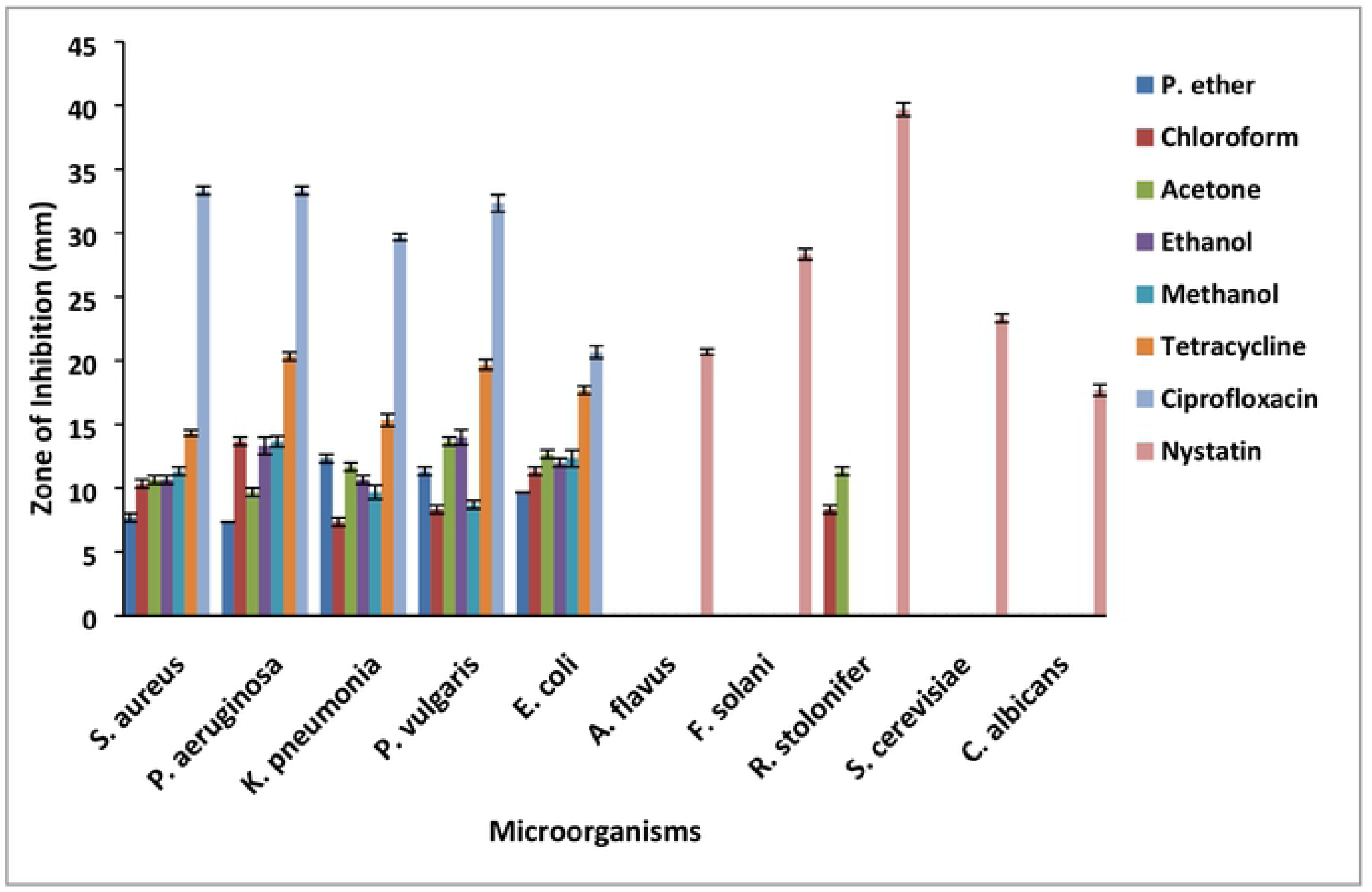
Antimicrobial activities of *S. xanthocarpum* leaves extract.

Among the different extracts of stem bark, the acetonic extract exhibited highest antimicrobial activity against *P. vulgaris* 14.67±0.33 mm (Table 3 and Fig 3). Methanol extract also showed promising activity against *S. aureus* (13±0.58 mm) followed by *P. aeruginosa* (12.67±0.33 mm). Along with that, no significant difference in zone of inhibition of methanolic extract (13±0.58 mm) and tetracycline (14.33±0.33 mm) against *S. aureus* was observed. The ethanol extract was found effective against *P. vulgaris* (12.67±0.33 mm). However, marginal inhibition was shown by *A. flavus (*8.67±0.33 mm) and *S. cerevisiae* (8.33±0.33 mm). Similar to fruit and leaf extracts, stem bark extract also showed insignificant antifungal potential. Among fungi, only *S. cerevisiae* showed significant zone of inhibition with petroleum ether and chloroform extracts. Present findings are similar to previous report of [15].

**Table 3:** Anti-microbial activity of *S. xanthocarpum* stem bark.

| Mean diameter of zones of inhibition (mm) ± Standard Error of Means (S.E.M) |  |  |  |  |  |  |  |  |  |
| --- | --- | --- | --- | --- | --- | --- | --- | --- | --- |
| S. No | Microorganisms | Petroleum ether | Chloroform | Acetone | Ethanol | Methanol | Tetracycline | Ciprofloxacin | Nystatin |
| 1 | *S. aureus* | 11.33±0.33e | 11.00±0.58e | 8.33±0.33f | 11.33±0.33e | 13±0.58d | 14.33±0.33d | 33.33±0.33a | --- |
| 2 | *P. aeruginosa* | 8.67±0.33f | 9.67±0.33f | 13.67±0.33d | 11.67±0.33e | 12.67±0.33e | 20.33±0.33c | 33.33±0.33a | --- |
| 3 | *K. pneumonia* | 10.33±0.33e | 8.33±0.33f | 10.33±0.33e | 10.67±0.33e | 11.33±0.33e | 15.33±0.33d | 29.67±0.33b | --- |
| 4 | *P. vulgaris* | 8.33±0.00f | 8.33±0.33f | 14.67±0.33d | 12.67±0.33e | 8.67±0.33 | 9.67±0.33 | 32.33±0.67a | --- |
| 5 | *E. coli* | 9.33±0.33f | 10.33±0.00e | 12.67±0.33e | 10.33±0.33e | 10.33±0.33e | 17.67±0.33d | 20.67±0.33c | --- |
| 6 | *A. flavus* | 0.00±0.00h | 0.00±0.00h | 0.00±0.00h | 8.67±0.33g | 0.00±0.00h | --- | --- | 20.67±0.33d |
| 7 | *F. solani* | 0.00±0.00h | 0.00±0.00h | 0.00±0.00h | 0.00±0.00h | 0.00±0.00h | --- | --- | 28.33±0.33b |
| 8 | *R. stolnifer* | 0.00±0.00h | 0.00±0.00h | 8.33±0.33g | 0.00±0.00h | 0.00±0.00h | --- | --- | 39.67±0.33a |
| 9 | *S. cerevisiae* | 11.33±0.33f | 10.33±0.33f | 0.00±0.00 | 8.33±0.33g | 0.00±0.00h | --- | --- | 23.33±0.33c |
| 10 | *C. albicans* | 0.00±0.00h | 0.00±0.00h | 0.00±0.00h | 0.00±0.00h | 0.00±0.00h | --- | --- | 17.67±0.33e |

Note: The means followed by same letters are non-significant

**Figure 3.**
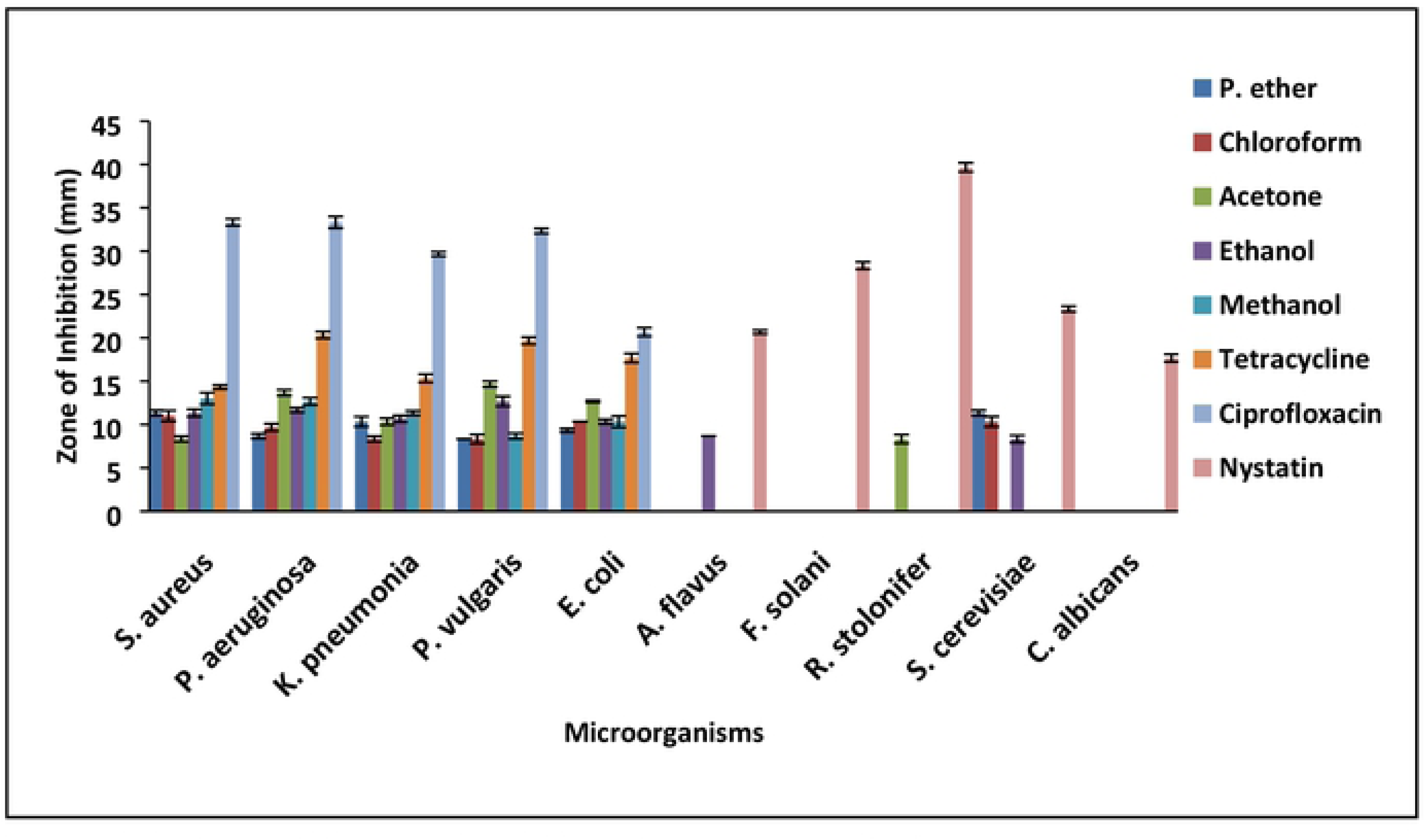
Antimicrobial activities of *S. xanthocarpum* stem bark extract.

The root bark extract of methanol showed maximum antimicrobial activity against *P. aeruginosa* and *P. vulgaris* (17.67±0.33 mm) each (Table 4 and Fig 4). The mild activity was observed against *K. pneumonia* (16.33±0.33 mm) and *S. aureus* (14.67±0.33 mm). Amongst all the tested fungi and yeasts, only *A. flavus* was found sensitive to methanol extract with zone of inhibition of 12.67±0.33 mm. All the other fungal strains were found to be resistant to root bark extract. Similar results were reported by [16]. Ethanol extract was also found to be active against all the tested bacteria followed by acetone, chloroform and petroleum ether extracts. *P. vulgaris* exhibited maximum sensitivity against methanol extract and *P. aeruginosa* against methanol and chloroform extract. The study of [17] also suggested that gram negative bacteria are more sensitive to extract than gram positive bacteria. Methanolic extract showed no activity against all tested fungi. These results are in good accordance with the previous studies which suggested that plant extracts are more effective against bacteria than fungi [18].

**Table 4:**
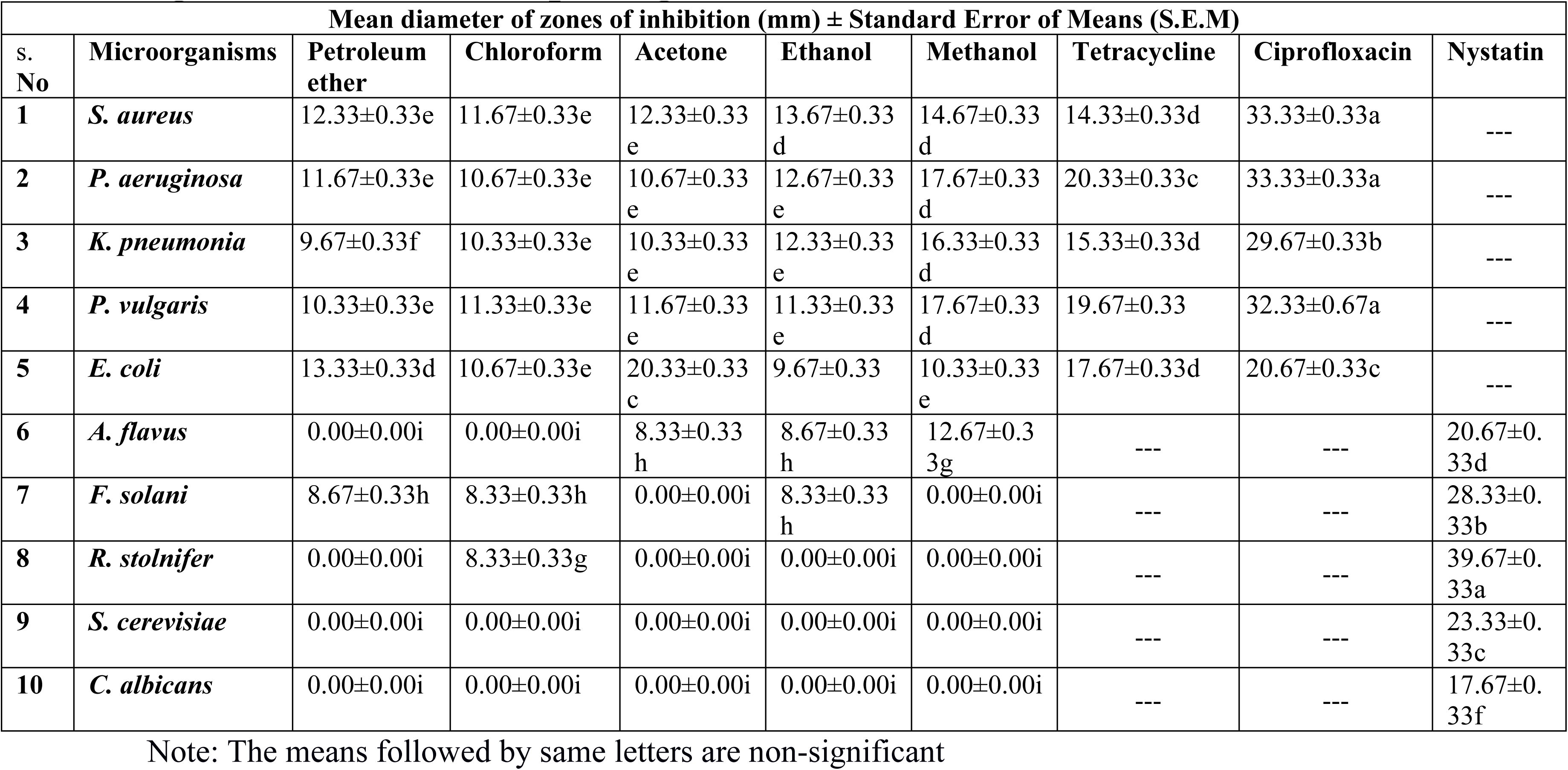
Anti-microbial activity of *S. xanthocarpum* root bark against tested human pathogens.

| Mean diameter of zones of inhibition (mm) ± Standard Error of Means (S.E.M) |  |  |  |  |  |  |  |  |  |
| --- | --- | --- | --- | --- | --- | --- | --- | --- | --- |
| S. No | Microorganisms | Petroleum ether | Chloroform | Acetone | Ethanol | Methanol | Tetracycline | Ciprofloxacin | Nystatin |
| 1 | *S. aureus* | 12.33±0.33e | 11.67±0.33e | 12.33±0.33e | 13.67±0.33d | 14.67±0.33d | 14.33±0.33d | 33.33±0.33a | --- |
| 2 | *P. aeruginosa* | 11.67±0.33e | 10.67±0.33e | 10.67±0.33e | 12.67±0.33e | 17.67±0.33d | 20.33±0.33c | 33.33±0.33a | --- |
| 3 | *K. pneumonia* | 9.67±0.33f | 10.33±0.33e | 10.33±0.33e | 12.33±0.33e | 16.33±0.33d | 15.33±0.33d | 29.67±0.33b | --- |
| 4 | *P. vulgaris* | 10.33±0.33e | 11.33±0.33e | 11.67±0.33e | 11.33±0.33e | 17.67±0.33d | 19.67±0.33 | 32.33±0.67a | --- |
| 5 | *E. coli* | 13.33±0.33d | 10.67±0.33e | 20.33±0.33c | 9.67±0.33 | 10.33±0.33e | 17.67±0.33d | 20.67±0.33c | --- |
| 6 | *A. flavus* | 0.00±0.00i | 0.00±0.00i | 8.33±0.33h | 8.67±0.33h | 12.67±0.33g | --- | --- | 20.67±0.33d |
| 7 | *F. solani* | 8.67±0.33h | 8.33±0.33h | 0.00±0.00i | 8.33±0.33h | 0.00±0.00i | --- | --- | 28.33±0.33b |
| 8 | *R. stolnifer* | 0.00±0.00i | 8.33±0.33g | 0.00±0.00i | 0.00±0.00i | 0.00±0.00i | --- | --- | 39.67±0.33a |
| 9 | *S. cerevisiae* | 0.00±0.00i | 0.00±0.00i | 0.00±0.00i | 0.00±0.00i | 0.00±0.00i | --- | --- | 23.33±0.33c |
| 10 | *C. albicans* | 0.00±0.00i | 0.00±0.00i | 0.00±0.00i | 0.00±0.00i | 0.00±0.00i | --- | --- | 17.67±0.33f |

Note: The means followed by same letters are non-significant

**Figure 4.**
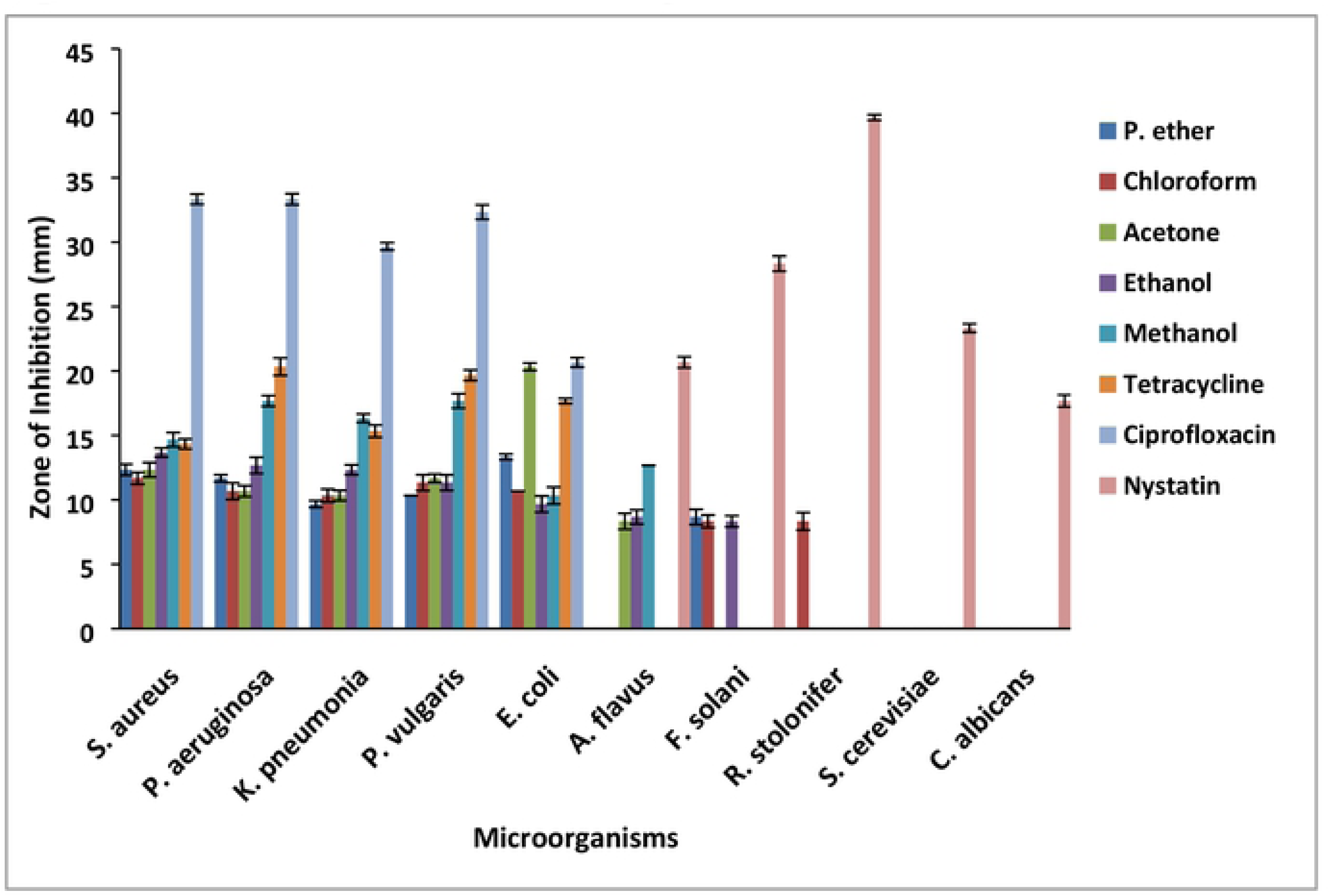
Antimicrobial activities of *S. xanthocarpum* root bark extract.

Phytochemical analysis of *S. xanthocarpum* revealed the presence of solasonine, solasomargine, sapogenin and solasodine that are responsible for their medicinal effects. Lipophilic flavonoids usually disrupt microbial cell membranes. Some phytochemicals form complexes with extracellular, soluble microbial proteins which bind to the microbial cell wall resulting in the dissolution of the cell wall [19]. It has been suggested that majority of phenolic compounds could be extracted with methanol extract except those phenolics which are bound to the insoluble carbohydrates and proteins [20]. Comparison with pertinent data from literature indicates that, according to the methodology adopted in studies on antimicrobial activity, the most diverse results can be obtained. Plant extracts have shown inhibitory effects on the growth of the bacteria studied, although of distinct forms. It is therefore recommended that the nature and the number of the active antibacterial principles involved in each plant extract should be studied in detail.

### 3.2 Antioxidant activities of different parts of *S. xanthocarpum*

The results of antioxidant activity revealed that highest IC_50_ (0.459242) was obtained ethanol extract of fruit and least with chloroform (0.842105) (Figure 5). When leaf extracts were investigated for their antioxidant activity, ethanol extract showed highest antioxidant potential with IC_50_ value 0.34188 and petroleum ether exhibited least antioxidant potential with IC_50_ value 0.672269. The methanol extract of stem bark showed highest antioxidant activity (0.323102) and acetone extract exhibited least activity (0.560224). Root bark demonstrated substantial antioxidant potential in chloroform (0.416667), ethanol (0.489596) and methanol (0.695652). Chloroform extracts of root bark revealed highest antioxidant activity. In this research work, methanol extract was found to possess higher antioxidant activity because the polar solvents allow extraction of all the phenolic or bioactive compounds from plants as compared to other solvents [21].

**Figure 5.**
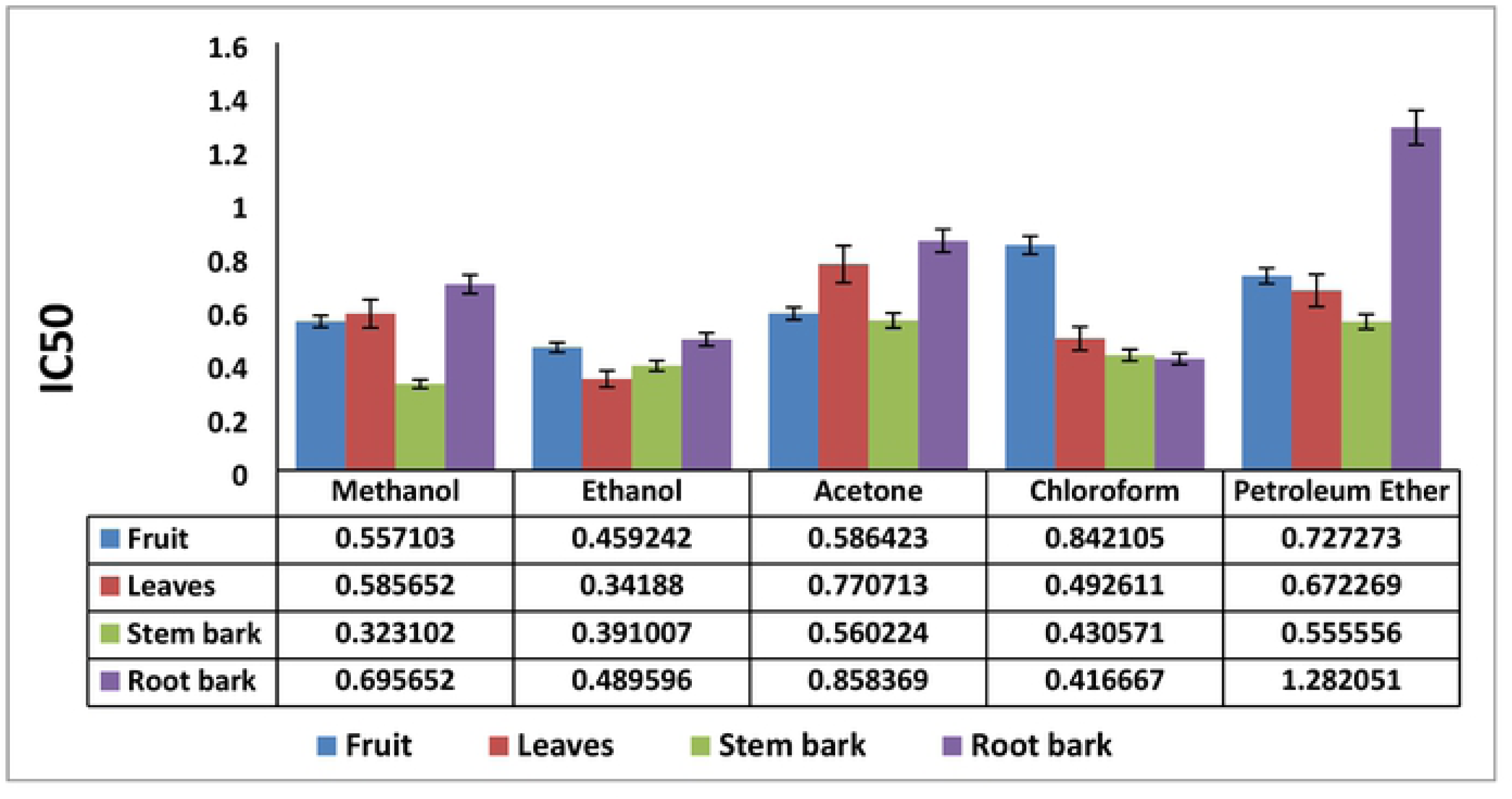
IC_50_ values of the four parts of *S. xanthocarpum* (fruits, leaves, stem bark and root bark).

### Phenolic Content of Different Parts of *S. xanthocarpum*

It is well established that phenolics and flavonoids are the important antioxidant substances that are obtained from most natural plants. Total phenolic content of methanolic extract of *S. xanthocarpum* (fruits, leaves, stem bark and root bark) are presented in Table 5 and Figure 6. Phenolics and flavonoids are strongly associated with significant reduction in cancer risk that are usually acquired from medicinal plants [22]. The phenolic content ranges from 12.3541 to 23.2942 Pmol GA/ug in different parts. The highest phenolic content was obtained from root bark extract (23.2942 Pmol GA/ug) and lowest in fruit extract (12.3541 Pmol GA/ug).

**Table 5:** Total phenolic content of *S. xanthocarpum* (fruits, leaves, stem bark and root bark) using Folin-Ciocalteu reagent.

| S. No. | Name of Plant | Part used | Conc. Used | Absorbance | (Pmol GA/ ug) |
| --- | --- | --- | --- | --- | --- |
| 1 | *S. xanthocarpum* | Fruits | 2.5 uL | 41.60 | 12.3541±1.73c |
| 2 | *S. xanthocarpum* | Leaves | 2.5 uL | 38.32 | 14.7065±2.32bc |
| 3 | *S. xanthocarpum* | Stem bark | 2.5 uL | 75.60 | 21.1082±2.88ab |
| 4 | *S. xanthocarpum* | Root bark | 2.5 uL | 80.50 | 23.2942±1.33a |

**Figure 6.**
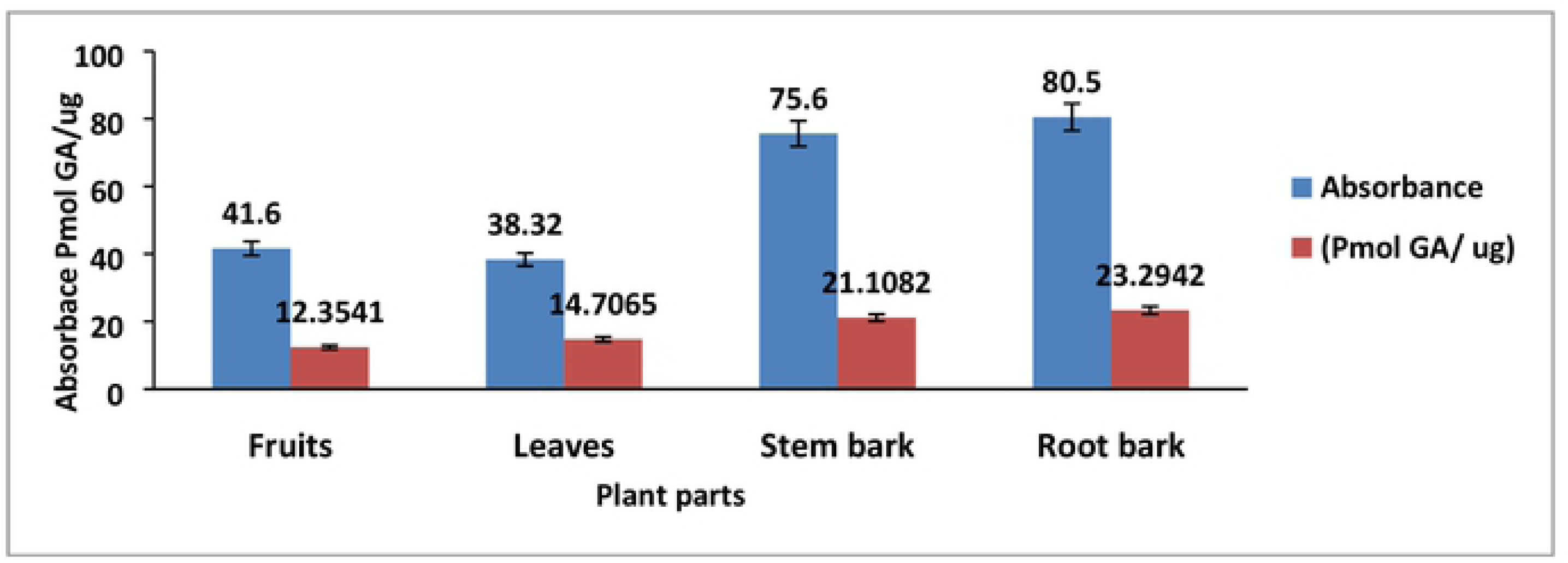
Total phenolic content in fruits, leaves, stem bark and root bark of *S. xanthocarpum*.

### Flavonoid Content in Different Parts of *S. xanthocarpum*

Flavonoid content in methanolic extracts of four parts (fruits, leaves, stem bark and root bark) of *S. xanthocarpum* are tabulated in table 6 and Figure 7. The highest flavonoid content was obtained from root bark extract (17.8480 Pmol GA/ug) and lowest in fruit extract (2.4806 Pmol GA/ug). Present results are in accordance with [23] that suggested cancer prevention agents belonging to a plant bioactive phytochemicals adjacent to phenolics, flavonoidsto be employed as additives against cancer.

**Table 6:** Total flavonoids contents of *S. xanthocarpum* (fruits, leaves, stem bark and root bark).

| S. No. | Name of Plant | Part used | Conc. used | Absorbance | (ug Rutin/ ug) |
| --- | --- | --- | --- | --- | --- |
| 1 | *S. xanthocarpum* | Fruits | 10 uL | 0.182 | 8.3341±1.73b |
| 2 | *S. xanthocarpum* | Leaves | 10 uL | 0.083 | 2.4806±0.59c |
| 3 | *S. xanthocarpum* | Stem bark | 10 uL | 0.334 | 15.3364±2.31a |
| 4 | *S. xanthocarpum* | Root bark | 10 uL | 0.304 | 17.8480±1.75a |

**Figure 7.**
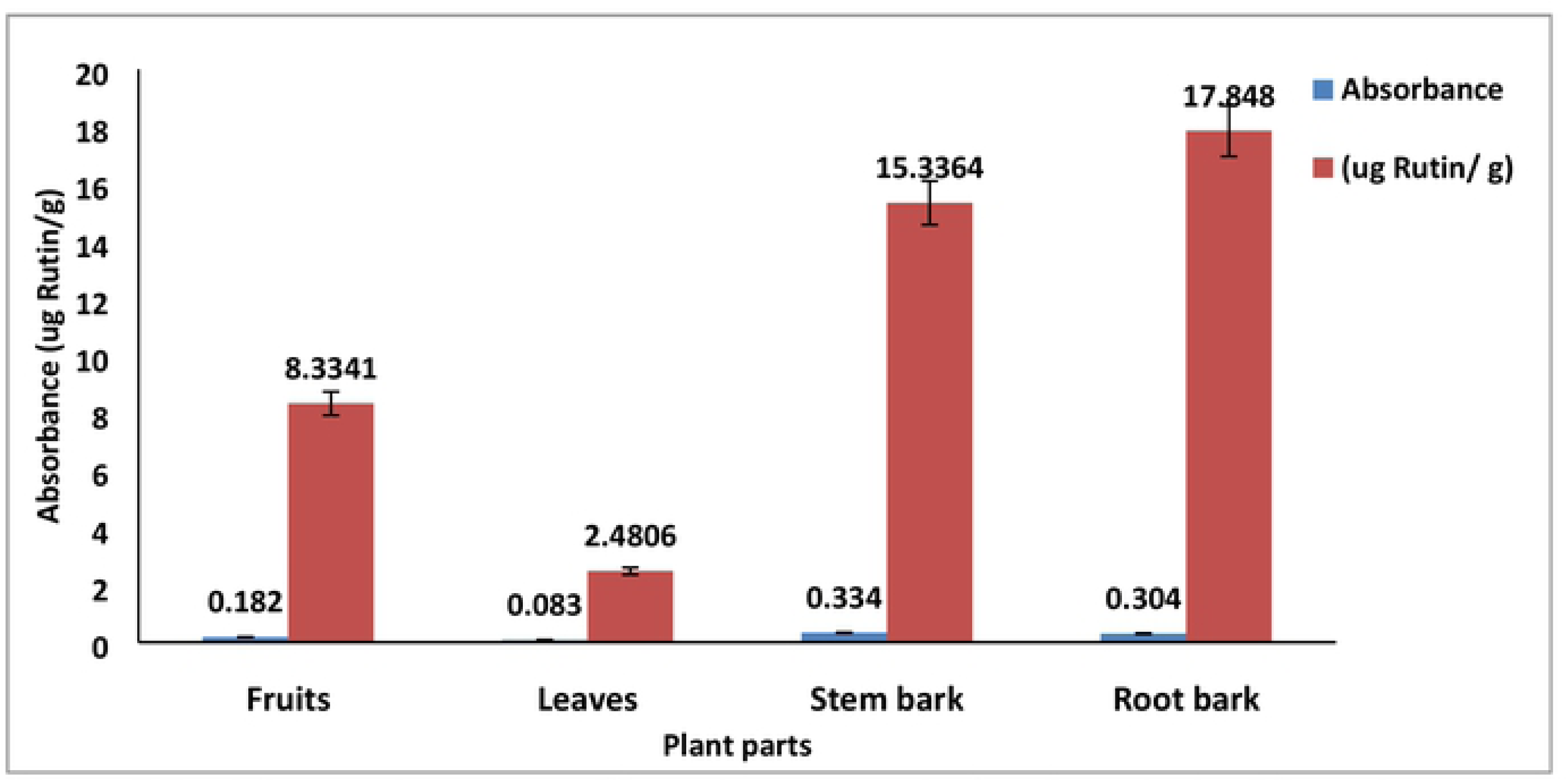
Total flavonoids contents of fruits, leaves, stem bark and root bark of *S. xanthocarpum*.

### Estimation of Metal Contaminations of *S. xanthocarpum*

The estimation of contamination of six metals (Cd, Cr, Cu, Mn, Mg and Zn) in four parts of *Solanum xanthocarpum* extracts was carried out by Atomic Absorption Spectrophotometry (AAS). The wave length to measure the concentrations of each metal in each extract and the mean concentration of each metal in each extract (in mg L^−1^) are given in Table 7 and Figure 8. The results of metal contamination were compared with the standard permissible limits (SPLs) by the FAO / WHO to evaluate their potential impact on the health. The Standard permissible limits (SPLs) of metal contamination (in dietary and medicinal species) are: Cd = 0.06 mg L-1, Cr = 0.05 mg L-1, Cu = 0.1 mg L-1, Mn = 0.26 mg L-1, Mg = 50 mg L-1 and Zn = 5.0 mg L-1 [24]. Present results showed that Cd concentration in stem bark and root bark, Cr concentration in leaves, stem bark and root bark, Mn concentration in leaves, Cu and Mg concentration in all parts of *S. xanthocarpum* (fruits, leaves, stem bark and root bark) are above the standard permissible limits. However, concentration of Zn in all parts is below the standard permissible limits. It was found that higher Cd concentration in stem bark and root bark of *S. xanthocarpum* was the major contributing reason to the potential health risk. Similarly, contamination of Cd has been reported in grape, spinach and celery [25, 26]. Therefore, these results should be considered as important guidelines for the producers, traders, consumers and regulatory authorities. Unfortunately, no regulatory practices exist for the assessment of metal contamination in the fruits of *S. xanthocarpum* which are extensively consumed by local population. The amount of Cu is critically above the standard permissible limits which indicate the high concentration of the Cu in these selected areas that ultimately accumulate in the flora, fruits and vegetables of this region.

**Table 7.** Metal concentration in *Solanum xanthocarpum* for the estimation of metal contamination (mg/L).

| # | Part Used | Cd | Cr | Cu | Mn | Mg | Zn |
| --- | --- | --- | --- | --- | --- | --- | --- |
| 1 | **Fruits** | 0.041±0.33<br>3h | 0.056±0.77<br>4f | 0.39±0.333<br>e | 0.196±0.77<br>4h | 98.4±0.152<br>c | 0.019±<br>0.819e |
| 2 | **Leaves** | 0.052±0.77<br>4g | 0.131±0.20<br>1e | 2.418±0.81<br>9c | 0.574±0.33<br>3d | 111.6±0.03<br>3b | 0.294±0.81<br>9e |
| 3 | **Stem bark** | 0.279±<br>0.819b | 0.193±0.77<br>4d | 3.106±<br>0.774b | 0.241±0.77<br>4f | 128±<br>0.881a | 0.318±<br>0.774e |
| 4 | **Root bark** | 3.675±0.52<br>7a | 0.211±0.77<br>4c | 3.731±<br>0.774a | 0.219±0.77<br>4g | 76± 0.577d | 0.394±0.77<br>4e |

**Figure 8.**
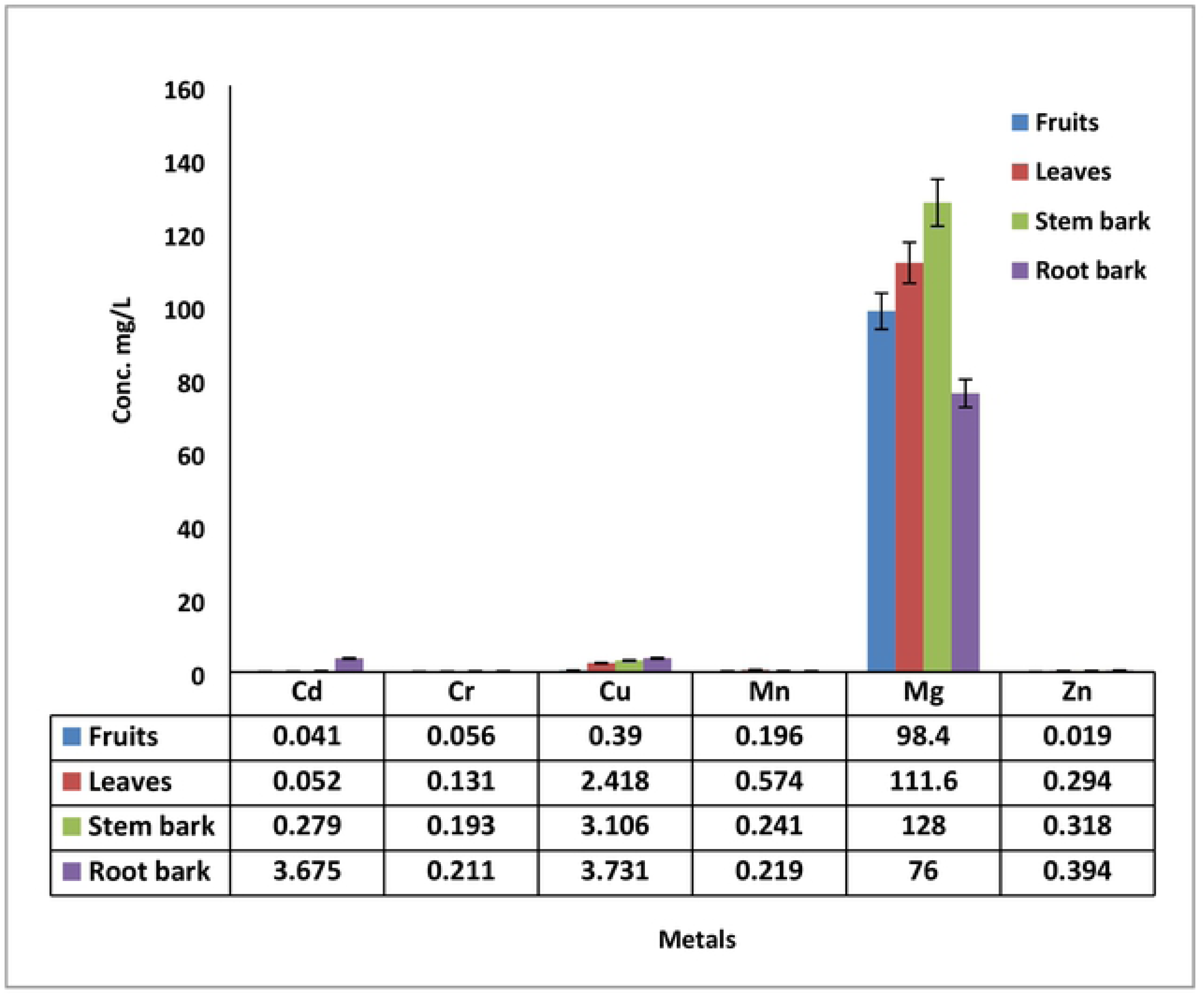
Metal contaminations in four parts of *S. xanthocarpum* (fruits, leaves, stem bark and root bark).

**Figure 9.**
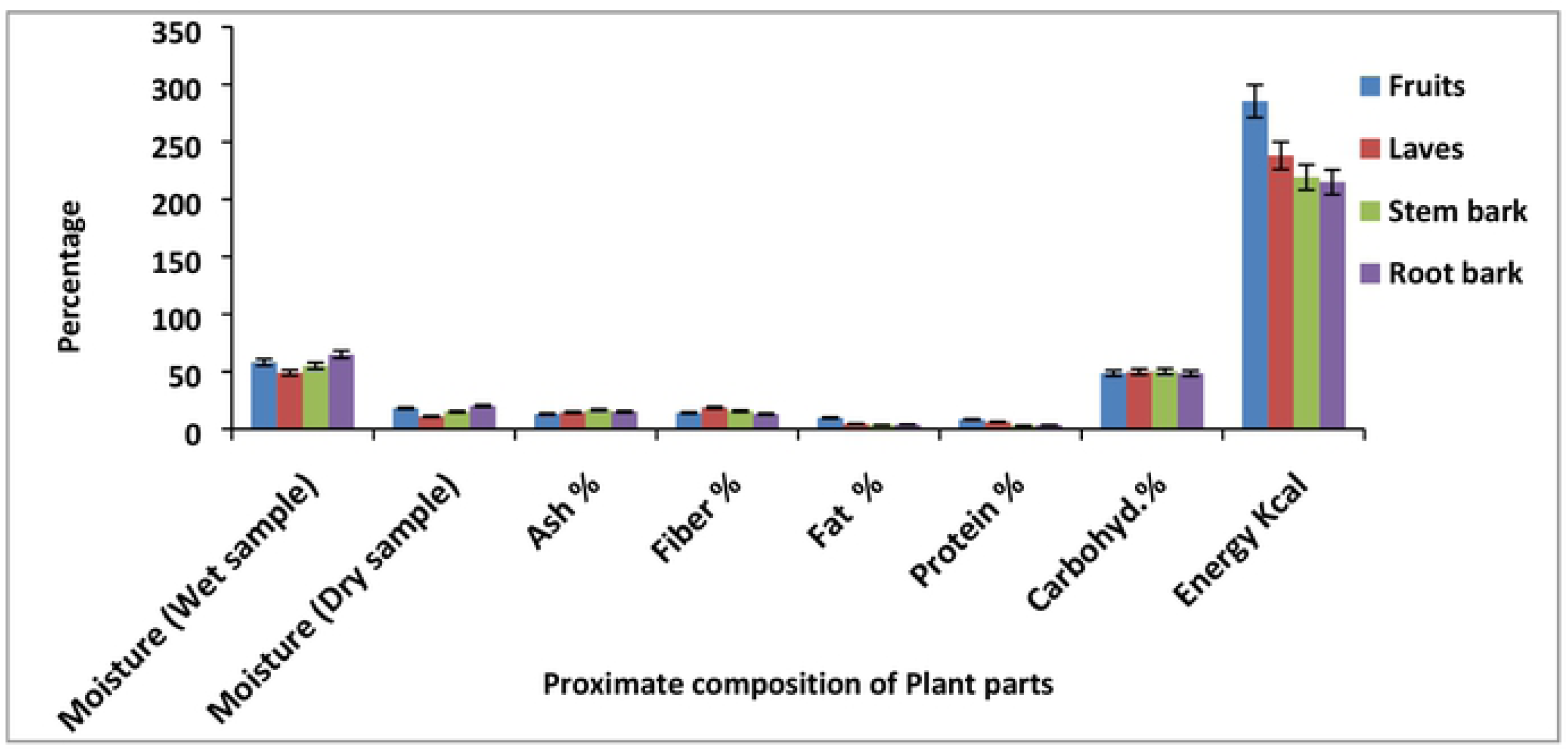
Proximate composition of *S. xanthocarpum* (fruits, leaves, stem bark and root bark).

In the present study, nutritional potential of *S. xanthocarpum* in Table 8 and Figure 9 was estimated to assess their prospective impact on the health. High moisture content (wet/dry) of all tested parts of *S. xanthocarpum*; fruits (58±0.577/18±0.577%), leaves (49±0.333/11±0.333%), stem bark (55±0.881/15±0.333%) and root bark (65±0.577/20±0.333%) was observed. Moisture contents determine the energy values in plants [27]. The highest moisture content (65%) of the root bark showed that selected plant is a good source of water for the healthier management of human body [28]. The moderate amount of ash content in all parts of *S. xanthocarpum* (13±0.577 to 16.50±0.33%) provides a measure of total amount of mineral matter in a plant and findings are similar to the values reported by [29] for some commonly consumed leafy vegetables. Ash content indicate the soluble and insoluble matter in the tested sample [30]. Crude fiber of *S. xanthocarpum* was found in fruits 14±0.333%, leaves 18.50±0.333%, 15.50±0.577% and root bark 13±0.577%. Crude fiber content of tested plant could play a significant role in intestinal bowel movement [31, 32]. Consuming vegetables in diet protect against cancer, lowers the cholesterol level and manages digestive disorders [33]. Comparatively lower fat content in all plants parts of *S. xanthocarpum* (3.42±0.037 – 9.79±0.065%) was observed. The low content of fat was reported for some leafy vegetables as plants are considered as poor sources of fats [34, 35]. Therefore, increased consumption of plants would naturally lower fat intake [36]. The protein content of the *S. xanthocarpum* tested parts was found to be moderate (2.93±0.015 – 8.32±0.105%). Protein is potential source for various functions of body such as building block of body development, fluid maintenance, enzymes, hormones formation and sustaining immunity [37]. The carbohydrate content in stem bark, fruit, root bark and leaves was found as 50.07±0.012%, 48.68±0.0584, 48.49± 0.77% and 49.74± 0.074% respectively. Comparatively high carbohydrate content in the stem bark confirm it as an essential source to supply energy required for the body [37, 38, 39]. High, moderate and low energy content of *S. xanthocarpum* was observed in fruits (285.455±0.527 Kcal), leaves (238.013±0.055 Kcal), stem bark (219.190±0.819 Kcal) and root bark (214.983± 0.819 Kcal). Almost similar results were also reported by [40].

**Table 8:** Proximate composition of *S. xanthocarpum* (fruits, leaves, stem bark and root bark).

| Nutritional values of *S. xanthocarpum* are expressed as mean ± Standard Error of three parallel replicates | | | | | | | | | |
| --- | --- | --- | --- | --- | --- | --- | --- | --- | --- |
| S. No | Plant's Part Used | Moisture (Wet sample) | Moisture (Dry sample) | Ash % | Fiber % | Fat % | Protein % | Carbohydrates % | Energy Kcal |
| 1 | Fruits | 58±0.577b | 18±0.577e | 13±0.577e | 14±0.333de | 9.79±0.065a | 8.32±0.105b | 48.68±0.058g | 285.455±0.527a |
| 2 | Laves | 49±0.333d | 11±0.333g | 14.50±0.333de | 18.50±0.333a | 4.53±0.157d | 6.26±0.017c | 49.74±0.074f | 238.013±0.055b |
| 3 | Stem bark | 55±0.881c | 15±0.333f | 16.50±0.333b | 15.50±0.577bc | 3.42±0.037e | 2.93±0.015f | 50.07±0.012e | 219.190±0.819c |
| 4 | Root bark | 65±0.577a | 20±0.333e | 15±0.333cd | 13±0.577e | 3.91±0.275e | 3.51±0.031e | 48.49±0.77h | 214.983±0.819d |

## Conclusion

It is concluded that *S. xanthocarpum* (fruits, leaves, stem bark and root bark) can be considered as potential source of anti-microbial, anti-oxidant compounds. Phenolics and flavonoids found in different parts of *S. xanthocarpum* can be used as food supplement and in pharmaceutical industry. Present investigations concluded that dietary and medicinal plant *S. xanthocarpum* should be analyzed for metal contamination before use. Proximate analysis showed that the *S. xanthocarpum* could be a good source of carbohydrate, moisture and energy, which may therefore justify both its nutritional and ethnomedicinal benefits to human health.

